# Virally induced CRISPR/Cas9-based knock-in of fluorescent albumin allows long-term visualization of cerebral circulation in infant and adult mice

**DOI:** 10.1101/2023.07.10.548084

**Authors:** Marta Vittani, Philip Aleksander Gade Knak, Masahiro Fukuda, Masaki Nagao, Xiaowen Wang, Celia Kjaerby, Ayumu Konno, Hirokazu Hirai, Maiken Nedergaard, Hajime Hirase

## Abstract

Albumin, a protein produced by liver hepatocytes, represents the most abundant protein in blood plasma. We have previously engineered a liver-targeting adeno-associated viral vector (AAV) that expresses fluorescent protein-tagged albumin to visualize blood plasma in mice. While this approach is versatile for imaging in adult mice, transgene expression vanishes when AAV is administered in neonates due to dilution of the episomal AAV genome in the rapidly growing liver. Here, we use CRISPR/Cas9 genome editing to insert the fluorescent protein mNeonGreen (mNG) gene into the albumin (Alb) locus of hepatocytes to produce fluorescently labeled albumin (Alb-mNG). We constructed a CRISPR AAV that includes ∼1 kb homologous arms around Alb exon 14 to express Alb-mNG. Subcutaneous injection of this AAV with AAV-CMV-Cas9 in postnatal day 3 mice resulted in two-photon visualization of the cerebral cortex vasculature within ten days. The expression levels of Alb-mNG were persistent for at least three months and were so robust that vasomotion and capillary blood flow could be assessed transcranially in early postnatal mice. This knock-in approach provides powerful means for micro- and macroscopic imaging of cerebral vascular dynamics in postnatal and adult mice.

## Introduction

Fluorescent labeling of blood plasma has been a popular method for imaging of cerebral microcirculation in the past few decades [1, 2]. Plasma labeling is typically performed by intravenous injection of fluorescent tracers (e.g., Texas Red, Fluorescein Isothiocyanate) conjugated to large dextran (size 80 to 2,000 kDa), whereby the vasculature and the flow therein can be visualized after a few seconds. In combination with two-photon microscopy [3], fluorescent plasma tracers have enabled investigation of cerebral vessels in normal and disease model rodents [4–9]. Convenient as these tracers are, major limitations include tracer leakage within few hours from administration, stress of repeated injections for longer experiments and introduction of biases on blood flow due to a higher blood viscosity.

To overcome these limitations, we have recently established a method by which fluorescent protein-tagged albumin is expressed following single-shot systemic administration of an adeno-associated viral vector (AAV) [10]. Albumin, a protein produced by liver hepatocytes, represents 50–60% of plasma proteins and exists at concentrations of 300–500 µM in adult rodents [11]. Albumin is negatively charged at physiological pH and is the sole plasma protein devoid of carbohydrate chains (i.e., non-glycoprotein), making it difficult to re-enter the cells once secreted [12]. We previously expressed a recombinant albumin protein with C-terminus fusion of mNeonGreen (Alb-mNG) in the liver by AAV carrying the hepatocyte-selective P3 transthyretin promoter [13]. We were able to image cerebral circulation two weeks after AAV administration for over four months, enabling investigation of longitudinal vascular function alterations at various spatial and temporal scales [10].

While AAVs can induce gene expression for months by retaining episomal circularized genomic components [14], the expression attenuates with cell turnover and division, since the viral genome is not integrated into the host cell’s genome. The liver grows rapidly between postnatal weeks two and eight to increase its weight by several fold in mice [15]. Accordingly, liver transgene expression vanishes in adults when AAV is administered in infants, whereas expression lasts for over four months with adult AAV administration [16].

Here, we introduced CRISPR/Cas9-based knock-in of mNG into the albumin locus by systemic AAV injection to produce fluorescent protein-tagged albumin (Fig. 1a). Since albumin is constantly produced with a half-life of 1.2–1.6 days in mice [17–19], we took advantage of the accessible chromatin structure for genome editing. We designed a CRISPR AAV to insert mNG into exon 14 of the mouse albumin genome to express mNG-fused albumin (Alb-mNG). The AAV vector contained a U6 promoter-driven sgRNA cassette targeting at the immediate downstream of *Alb* exon 14 and a DNA sequence that adds a *mNG* coding sequence to the 5’-end of *Alb* with ∼1000 bp homologous arms in both directions (Fig. 1b). The resulting AAV9-U6-AlbEx14-mNG was co-injected with AAV9-hCMV-HA-SpCas9 (Fig. 1c) subcutaneously in postnatal day 3 (PN3) mice. Successful production of Alb-mNG was confirmed by *ex vivo* blood sample fluorescence for at least 15 weeks post-injection. The cerebral cortex vasculature was visualized transcranially within 7–10 days from injection by two-photon imaging. Plasma levels of mNG and liver transcript levels of mNG over total albumin were measured to further validate the model. Overall, we demonstrate that AAV-mediated genome-editing represents a powerful strategy for the visualization of blood flow by fluorescently labeled albumin and enables investigation of murine vasculature from neonatal stages.

**Fig. 1.**
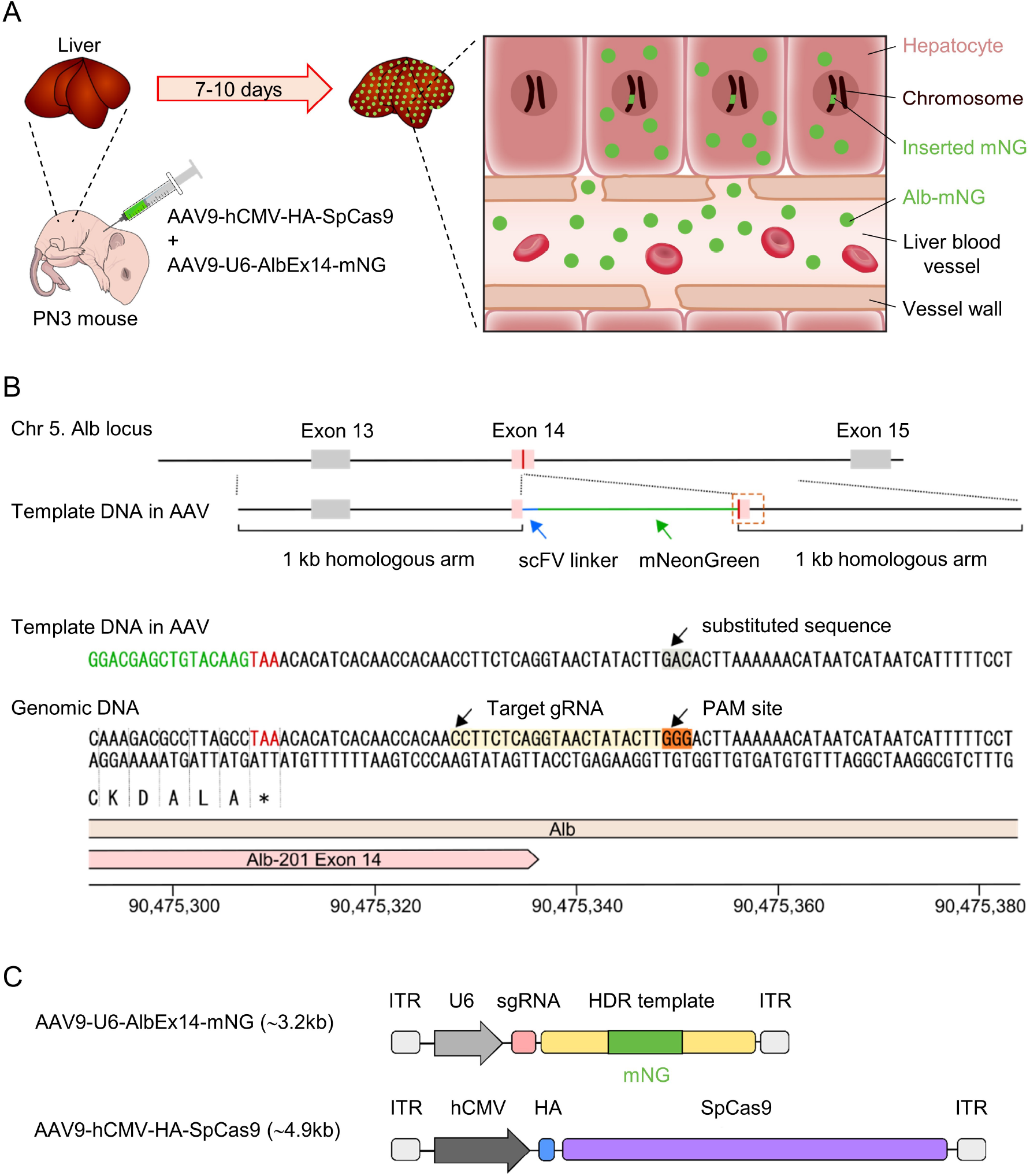
Production of Alb-mNG by genome-edited hepatocytes. (A) Experimental approach for genome editing of hepatocytes to produce albumin tagged with the fluorescent protein mNeonGreen (Alb-mNG). An AAV mixture carrying CRISPR (AAV9-U6-AlbEx14-mNG) and Cas9 components (AAV9-hCMV-HA-SpCas9) is administered subcutaneously to a PN3 mouse. Within 7–10 days from injection, mNG is inserted to the albumin locus of hepatocytes’ genome, leading to persistent production of Alb-mNG and its secretion into the blood stream. (B) Schematic design and sequences of the template DNA in CRISPR AAV (AAV9-U6-AlbEx14-mNG) and of the targeted albumin locus on chromosome 5. CRISPR AAV contains a U6 promoter-driven sgRNA cassette targeting at the immediate downstream of Alb exon 14 on chromosome 5 and a template DNA sequence that adds a mNG coding sequence to the 5’-end of Alb with ∼1000 bp homologous arms in both directions. The sgRNA is coded by 5’-CTTCTCAGGTAACTATACTT-3’ and targets the intron protospacer adjacent motif (PAM) site GGG after Alb exon 14. In the template DNA, the stop codon of Alb (TAA) is replaced with the scFV linker sequence followed by mNG-stop. In addition, the intronic PAM site for AlbEx14-mNG is substituted with GAC. The genomic map is from the Genome Reference Consortium Mouse Build 39 (GRCm39). (C) Schematic design of AAV9-U6-AlbEx14-mNG and AAV9-hCMV-HA-SpCas9.

## Materials

### 1. AAV preparation

AAV for knocking in *mNG* to the *Alb* locus by CRISPR/Cas9 gene editing was made following Nishiyama et al. [20]. The guide RNA was designed according to the sgRNA synthesis protocol in Mali et al. [21] using the B6j mouse reference genome GRCm39. An intron PAM site after *Alb* exon 14 on chromosome 5 at position 90475346 containing GGG on the positive strand was targeted. The construct, which we named AlbEx14-mNG, contained a sgRNA coded by 5’-CTTCTCAGGTAACTATACTT-3’ and a targeting sequence for homologous recombination. The targeting sequence consisted of 1000 bp homology arms prior and posterior to the PAM site, whereby the stop codon of Alb was replaced with the scFV linker sequence followed by mNG-stop. In addition, the intronic PAM site for AlbEx14-mNG was replaced with GAC. AlbEx14-mNG was assembled with the U6 promoter and sgRNA scaffold and subcloned into an AAV plasmid (Addgene 105550, pAAV-GFAP-Cre, gift from James M. Wilson) via NotI. This plasmid has been deposited at Addgene as pAAV-U6-AlbEx14-mNG (#201780). The AAV was purified with the AAV9 capsid (AAV9-U6-AlbEx14-mNG) by the Viral Core Facility at Charité – Universitätsmedizin Berlin yielding ∼1×10^13^ vg/mL (*see* Notes 1.1 and 1.2). The AAV for Streptococcus pyogenes Cas9 expression was obtained from the Viral Vector Facility, Institute of Pharmacology and Toxicology, University of Zurich (purified by VVF as v723-9; AAV9-hCMV-HA-SpCas9, Addgene 106431, gift from Juan Belmonte [22]).

### 2. Mice for AAV injection

Infant and adult C57BL/6JRj mice of both sexes were used. Two-week pregnant females were ordered from Janvier (*see* Note 1.3). Infant mice were housed with the mother until weaning at 21 days. Mice were housed in normal 12-h light/12-h dark cycle (lights on: 7am) with food and water *ad libitum*. The procedures involving animal care, surgery, *in vivo* imaging, and sample preparation were approved by the local research ethics committee (Department of Experimental Medicine, University of Copenhagen) and conducted in accordance with the Danish Animal Experiments Inspectorate.

### 3. AAV administration

AAV9-U6-AlbEx14-mNG, AAV9-hCMV-HA-SpCas9, and sterile saline were mixed to reach a final injection volume of 50 µL using a vortex mixer. Subcutaneous injection was chosen as administration route (*see* Note 1.4). The AAV mixture was loaded into a BD Micro-Fine Demi 0.3mL U-100 Insulin syringe w. 30G needle (BD, #3079125). PN3 mice were briefly anesthetized using isoflurane prior to injection and their body temperature was maintained using a heating pad during recovery from anesthesia. Once fully recovered, injected mice were placed back with the dam.

### 4. *Ex vivo* macro fluorescence imaging

Awake animals were sampled at PN7 at the earliest and at 15 weeks of age at the latest. Needles (30G–25G) were used to puncture the tail tip of the animals. Blood was collected from the punctured site with borosilicate glass capillaries (1B100F-4 or 1B150F-4, WPI). Fluorescence of the blood samples was examined by a macroscope (Leica M205 FA) equipped with an X-Cite 200Dc light source, digital camera (C11440 Orca-flash 4.0, Hamamatsu). Filter sets ET GFP LP (excitation 480/40, emission 510LP, 10447407, Leica) were used to image green. Images were acquired using Leica Application Suite X software (version 2.0.0.14332.2) with the following imaging settings: zoom of 2X, exposure time of 1.5s.

### 5. Acute head-plating and two-photon imaging

PN10–PN15 mice were anesthetized with ketamine (50 mg/kg) - xylazine (5 mg/kg) and mounted on a stereotaxic frame. Their body temperature was maintained at 37 °C using a heating pad. The skull was exposed, and a metal head plate was attached to the skull using dental cement (Super Bond C&B, Sun Medical, Shiga, Japan). The anesthetized mouse was then head-fixed under a two-photon microscope and its skull surface was covered with hydrogel (e.g. ultrasound gel Dahlhausen #93.050.00.443) to maintain it hydrated and transparent. During the recording session (max 2 h long), body temperature was maintained using a heating pad and anesthesia depth was frequently assessed so that to re-dose with ketamine 50 mg/kg if necessary. Two-photon imaging was performed using a Bergamo microscope (Thorlabs) equipped with a resonant scanner, a Chameleon Ultra 2 laser (Coherent), an objective lens (XLPlan N; ×25 NA = 1.05; Olympus) and the primary dichroic mirror FF705-Di01-25×36 (Chroma). Emission light was separated by the secondary dichroic mirror (FF562-Di03, Semrock) with band-pass filters FF03-525/50 and FF01-607/70 (both Semrock). mNG was excited at 940 nm. Images were acquired using ThorImage Software Version 3.0. The laser power under the objective lens was measured by a power meter (Thorlabs) before imaging.

### 6. Plasma and liver collection

Deeply anesthetized mice were intraperitoneally injected with heparin (500U, LEO) 30 minutes before collection to prevent blood clotting. The heart was exposed, and the right atrium was excised. The blood was quickly collected with a syringe and transferred in 0.75 mL tubes containing 5 μL EDTA and 5 μL Halt protease and phosphatase inhibitor cocktail (100x, Thermo Scientific). Blood samples were centrifuged for 10 min at 2000 x g at 4°C. The supernatant, i.e., plasma, was collected and stored in aliquots at −80°C. From the same animals, liver was also collected immediately after blood and either flash-frozen or submerged into RNAprotect Tissue Reagent (Qiagen, #76104). Liver samples were stored at −80°C.

### 7. Plasma mNG quantification

Plasma samples of injected mice were diluted at 1:30 in PBS and their fluorescence was measured in duplicates using a SpectraMax iD3 plate reader (excitation 485 nm, emission 535 nm). Six serial dilutions with known mNG concentration ranging from 6.25 to 200 nM were created by reconstitution of mNG protein (Gene Universal, USA) in PBS. These were loaded on the same microplate as the samples in order to create a standard curve. The fluorescence values from the experimental samples were linearly fit to the standard curve and the dilution factor 30 was considered to get plasma mNG concentration of the injected mice.

### 8. Gene expression analysis by real-time quantitative PCR

RNA was extracted from liver tissues using RNeasy Protect Mini Kit (Qiagen, #74124). The total RNA extract was then quantified, and its quality was checked by agarose gel electrophoresis. RNA was reverse transcribed into complementary DNA (cDNA) using iScript cDNA Synthesis Kit (Bio-Rad, #1708891). Several primer pairs specifically targeting mNG and albumin were designed and tested with qualitative PCR. The following primer pairs were chosen: mNG FW 5’-TGGTGCAGGTCGAAGAAGAC-3’, mNG RV 5’-GCTTGCCATTTCCAGTGGTG-3’; Alb FW 5’-TTTCGCCGAGAAGCACACAA-3’ (targeting the junction between exons 1 and 2), Alb RV 5’-ACTGGGAAAAGGCAATCAGGA-3’ (targeting exon 3). SsoFast™ EvaGreen® Supermix 2X (Bio-Rad, #172-5204) was used for qPCR. For each tested gene, a standard curve was established by using a dilution series of the pooled cDNA samples to assess the reaction efficiency and determine the initial starting amount of the target template. For gene expression analysis, relative quantification was made with actin as reference gene: Actb FW 5’-AGTGTGACGTTGACATCCGTA-3’, Actb RV 5’-GCCAGAGCAGTAATCTCCTTCT-3’.

## Methods

### 1. Subcutaneous injection of AAV mixture carrying CRISPR and Cas9 components in PN3 mice

An AAV mixture was prepared by mixing AAV9-U6-AlbEx14-mNG at a dose of 2 × 10^11^ vg with AAV9-hCMV-HA-SpCas9 at a dose of 2 × 10^11^ vg (*see* Note 1.5). The final injection volume of 50 µL was reached by adding sterile saline (*see* Note 1.6). Prior to injection, the AAV mixture was vortexed (*see* Note 1.7) and collected with a 0.3mL insulin syringe with 30G needle (*see* Note 1.8). Each PN3 mouse was briefly anesthetized by isoflurane at high concentration (3-4%) and transferred on a heating pad in prone position (*see* Note 1.9). Its loose skin on the neck was gently pinched to allow insertion of the 30G needle and subcutaneous administration of the AAV mixture. After needle removal, the mouse was kept under observation for a few minutes until recovery from anesthesia before returning to its home cage (*see* Note 1.10).

### 2. *Ex vivo* validation by macro fluorescence imaging

Macro fluorescence imaging of blood samples in glass capillaries was used as a fast and reliable *ex vivo* method to assess expression strength in single animals before proceeding to more elaborate experiments (*see* Note 2.1). While gently holding the tail of an unanesthetized mouse, a needle (*see* Note 2.2) was used to puncture the tail tip to collect a few microliters (μL) of blood in a glass capillary from the punctured site (*see* Note 2.3). Fluorescence was then examined under a macroscope (*see* Note 2.4). The same imaging settings were used across different animals and studies to allow for comparison (*see* Note 2.5). For longitudinal assessment of plasma fluorescence, an un-injected (naïve) mouse was also sampled for control reference.

This method allowed us to examine the long-term expression of Alb-mNG. The fluorescence signal was already detectable 4 days after injection and progressively increased until peaking at 4–5 weeks post-injection (Fig. 2a–b). The expression lasted for over 15 weeks from injection (*see* Note 2.6). The gradual temporal profile of mNG expression is likely a reflection of sgRNA expression, Cas9 concentration, and transport of AAV genome to the nucleus, which are essential components of homology-directed repair. Of note, a simpler AAV-mediated CRISPR/Cas9 knockout based on non-homologous end-joining is reported to work in a similar time course [23]. Afterwards, AAVs are progressively degraded and Cas9 is attenuated due to the CMV promoter silencing [24], thus slowing down the formation of newly edited hepatocytes. Around 8 weeks of age, the liver reaches its full size [15], while maintaining a pool of knock-in hepatocytes constantly producing fluorescent albumin. Remarkably, the plasma mNG fluorescence at plateau (8–15 weeks) is comparable to samples from our previous work, in which microcirculation was visualized *in vivo* by promoter-driven overexpression of Alb-mNG by AAV [10].

**Fig. 2.**
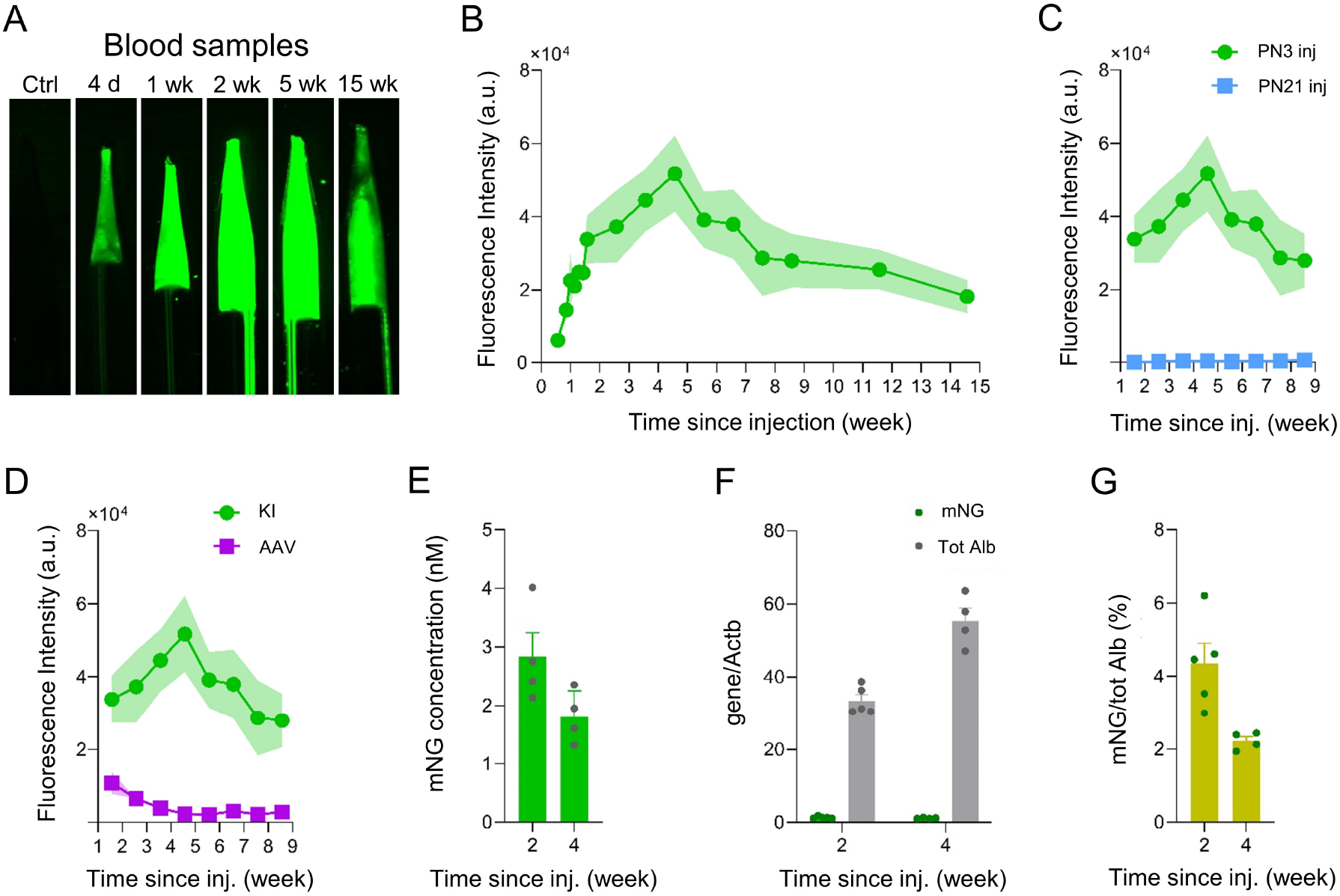
Quantification of blood Alb-mNG levels and estimation of knock-in efficiency. (A) Example of the fluorescence signals in blood samples collected on day 4 and weeks 1, 2, 5 and 15 from a mouse that received subcutaneous injection of AAV9-CMV-Cas9 and AAV9-U6-AlbEx14-mNG at PN3. (B) Alb-mNG fluorescence intensity in blood samples of mice injected with AAV9-CMV-Cas9 and AAV9-U6-AlbEx14-mNG at PN3 over a time course of 15 weeks (n = 2–5 mice). (C) Alb-mNG fluorescence intensity in blood samples of mice injected with AAV9-CMV-Cas9 and AAV9-U6-AlbEx14-mNG at either PN3 or PN21 over a time course of 8 weeks (n = 3 mice). (D) Alb-mNG fluorescence intensity in blood samples of mice injected with AAV9-CMV-Cas9 and AAV9-U6-AlbEx14-mNG (KI) or AAV8-P3-Alb-mNG (AAV) at PN3 over a time course of 8 weeks (n = 3 mice). (E) mNG concentration in plasma samples collected on weeks 2 and 4 from mice injected with AAV9-CMV-Cas9 and AAV9-U6-AlbEx14-mNG at PN3 (n = 4–6 mice). (F and G) Transcript levels of mNG or total albumin normalized on reference Actin in liver samples collected on weeks 2 and 4 from mice injected with AAV9-CMV-Cas9 and AAV9-U6-AlbEx14-mNG at PN3 (n = 5–6 mice). Percentage of mNG over total albumin transcript levels. Graphs show means ± SEM and individual values.

Injection of the same AAV mixture at PN3 versus PN21 (weaning) was compared up to 9 weeks from injection (Fig. 2c). The latter did not result in detectable fluorescence signal of Alb-mNG, most probably due to insufficient AAV dosage for the adult liver. Therefore, it is critical to inject at very early postnatal stages, such as PN3, for the tested dosage of 2 × 10^11^ vg (*see* Note 2.7).

We also compared injection of the knock-in AAV mixture with injection of AAV8-P3-Alb-mNG from our previous work [10] in PN3 mice (Fig. 2d). Despite some barely detectable fluorescent levels around 2 weeks post-injection, AAV8-P3-Alb-mNG did not produce consistent levels of fluorescent albumin. Indeed, as this AAV is not integrated in the genome and exists in the cytosol as an episome, it gets diluted in the rapidly growing liver until disappearance.

### 3. A possible application: transcranial imaging in PN10-15 mice

Mice in the postnatal period have fairly thin skulls, thus allowing for transcranial imaging. Since the mNG fluorescence in the blood in PN10–15 mice appeared comparable to samples from our previous work, in which microcirculation was visualized *in vivo* by overexpression of Alb-mNG by AAV [10], we sought to image cerebral microcirculation through the skull by a two-photon microscope. Mice were considered ready for transcranial imaging once the fluorescence signals of mNG from blood samples reached around 2×10^4^ a.u. (Fig. 2b) (*see* Note 3.1). This occurred around 1–2 weeks after injection (i.e., in PN10–PN15). Mice were anesthetized with ketamine-xylazine, head-plated, and imaged acutely. We captured volumetric cortical microvasculature images showing big superficial vessels and deeper capillary nets (Fig. 3a–b). High frame rate imaging (>160Hz) of a single capillary allowed us to follow the movement of single red blood cells through the skull (Fig. 3c). The intensity of the fluorescent signal was sufficient to record spontaneous vasomotion, allowing discernment of arterioles and venules (Fig. 3d–g).

**Fig. 3.**
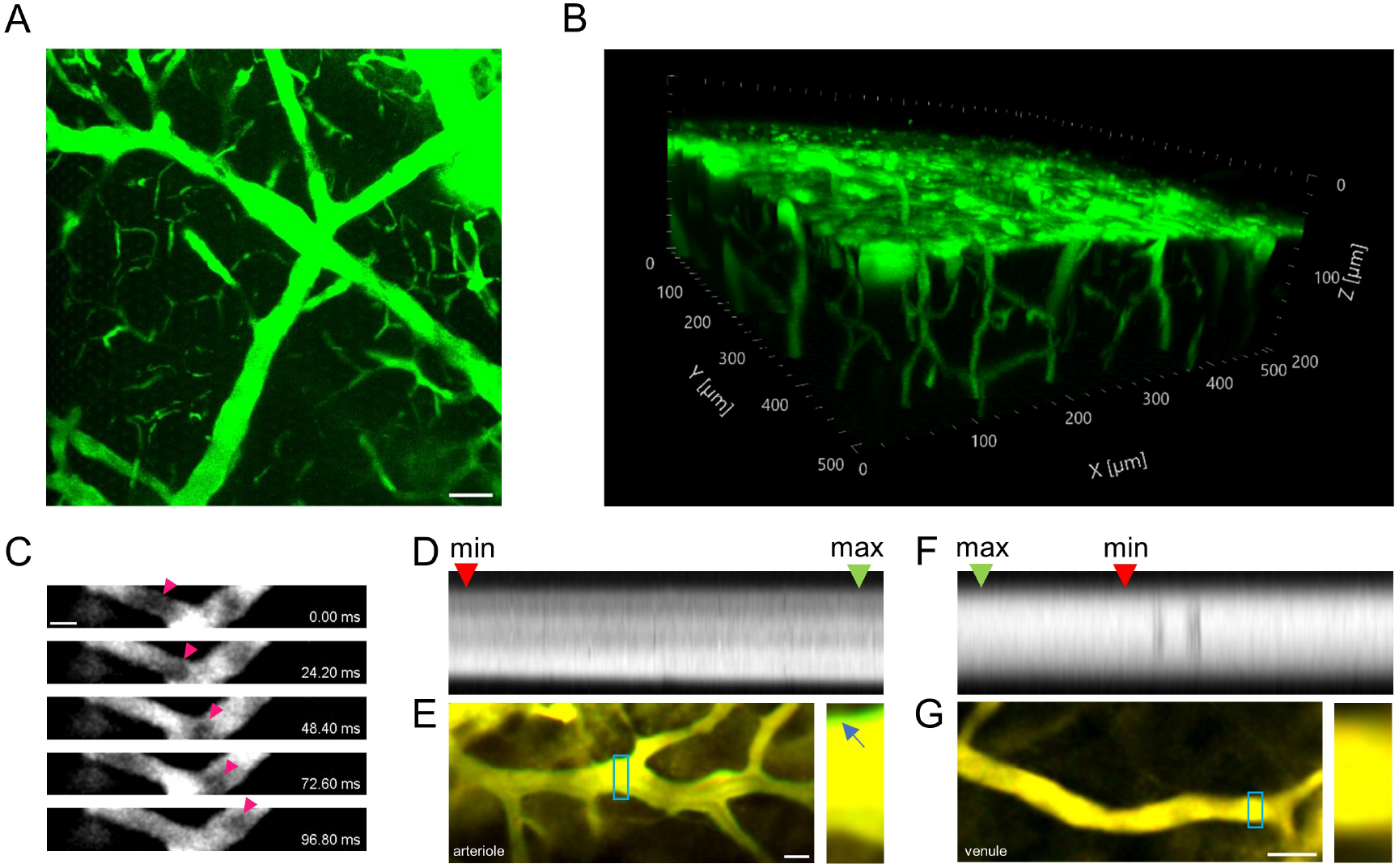
Transcranial imaging of Alb-mNG by two-photon microscopy within 10 days from injection. (A) Maximum intensity projection of a 130 µm z-stack showing cortical vasculature below the dural surface of a PN12 Alb-mNG-expressing mouse (post-injection 9 days). Scale bar, 50 μm. (B) Volumetric imaging of brain vasculature covering 200 µm with skull, meninges and cortex of a PN12 Alb-mNG-expressing mouse (post-injection 9 days). Scale bar, 514.02 μm x 514.02 μm x 200 μm. (C) High frame rate recordings of blood flow in a capillary of a PN12 Alb-mNG-expressing mouse (post-injection 9 days). Pink triangles indicate the flow of an example red blood cell. Scale bar, 5 μm. (D and E) Kymograph generated from ROI (blue square) of an arteriole in a PN12 Alb-mNG-expressing mouse (post-injection 9 days). Superimposed image of ROI with minimum arteriole diameter in red and maximum in green. Zoomed in area representing ROI (artery dilation in green indicated by the arrow). (F and G) Kymograph generated from ROI (blue square) of a venule in a PN13 Alb-mNG-expressing mouse (post-injection 10 days). Superimposed image of ROI with minimum vein diameter in red and maximum in green. Zoomed in area representing ROI (no vein dilation). Scale bars, 20 μm.

### 4. Post-mortem validation by biochemical assays

To further validate the model, plasma and liver tissues were collected from mice 2 and 4 weeks post-injection (*see* Note 4.1). Quantification of plasma mNG by comparing fluorescence with mNG standards indicated ∼2.8 μM and ∼1.8 μM plasma Alb-mNG at 2 and 4 weeks after injection, respectively (Fig. 2e). Genomic incorporation of *mNG* in the liver tissue, in which ∼70% is hepatocytes [25], was analyzed by qPCR for *mNG* and for *Alb*. Transcript levels of mNG were similar at the two time points (1.4 and 1.2, Fig. 2f), whereas total albumin mRNA levels almost doubled between 2 and 4 weeks post-injection (33.5 and 55.3, Fig. 2f). The knock-in efficiency — measured as the percentage of *mNG* transcript levels over total albumin transcript levels — was respectively 4.4% and 2.2% at 2 and 4 weeks post-injection (Fig. 2g).

## Notes

### Note 1: Subcutaneous injection of PN3 mice

Note 1.1: AAV serotype 9 does not target the liver specifically, but the liver is always strongly infected possibly owing to the large endothelial fenestrations (>100 nm) present in hepatic capillaries. Furthermore, this serotype has high tropism to hepatocytes [26, 27].

Note 1.2: It is essential to purify AAVs with a high titer (minimum ∼1×10^13^ vg/mL). This will allow administration of high AAV dosages despite the major volume limitation when injecting PN3 mice (*see* Note 1.4).

Note 1.3: An alternative is in-house breeding.

Note 1.4: The major limitation of injecting mice at PN3 is the limited injectable volume, due to the low weight of the animals. The maximum injectable volume varies according to the administration route: subcutaneous injection allows up to 50 µL, whereas intraperitoneal injection and intravenous injection via retro-orbital sinus only up to 20 µL [28]. Having to inject two different AAVs, each at high dosage, we preferred to maximize injection volume by choosing subcutaneous injection.

Note 1.5: The dosage was determined considering that an adult murine liver should contain an order of 10^8^ hepatocytes, given a liver mass of 1.5 g and a hepatocyte density of 1.35×10^8^ cells/g [15, 29]. While systemic AAV9 administration leads to infection in hepatocytes and other cells, Alb-mNG should principally be produced by hepatocytes due to their predominant production of albumin. Therefore, the selected dosage of 2 × 10^11^ vg should be sufficient for broad infection of hepatocytes in the growing liver.

Note 1.6: We recommend to always inject the maximal injectable volume of 50 µL and to always include saline in the mixture of high titer AAVs to lower its viscosity and make the injection smoother.

Note 1.7: Vortexing is essential for thorough mixing of the highly viscous AAV mixture.

Note 1.8: We recommend using a 0.3mL insulin syringe with a 30G needle to minimize loss of AAV mixture in the syringe dead space and to limit tissue damage at injection point.

Note 1.9: Anesthesia is optional. We recommend to briefly anesthetize PN3 mice with isoflurane to attenuate erratic movements, thus reducing the risk of tissue damage or failed injection when administering AAVs. When placed on the heating pad, the animal is no longer under isoflurane. However, the mouse is still unconscious and immobile since the injection takes only 10–20 seconds.

Note 1.10: When returning the pups to their home cage, we recommend to gently rub the pups with nesting materials from the same cage so to eliminate external odors and reduce the possibility of maternal cannibalism.

Note 1.11: It is known that neutralizing antibodies produced after a first exposure to AAV administration negatively affects the efficiency of subsequent AAVs injections [30]. We have not tested co-injection of other AAVs in this protocol. We note, however, that it is possible to target hepatocytes by AAV at a later time via intraperitoneal injection (unpublished data), but not via intravenous injection via the retro-orbital sinus [10].

### Note 2: *Ex vivo* macro fluorescence imaging

Note 2.1: We strongly recommend not skipping this step. It is a simple way to confirm successful Alb-mNG production before proceeding to more time-consuming and expensive procedures, such as head-plating and two-photon imaging.

Note 2.2: The needle size varies according to the age of the animal: 30G needles for pups until weaning, i.e., PN21; 27G or 25G needle for weaned mice.

Note 2.3: If the amount of blood exiting from the punctured site is not enough, gently press the tail from the bottom towards the tip to promote blood stream.

Note 2.4: The blood sample should be imaged immediately after collection to avoid extensive blood coagulation.

Note 2.5: Quantifying blood samples provides a way to compare levels of Alb-mNG in the same mouse at different time points and in different mice at multiple time points. There are two kinds of variabilities that need to be considered. The first one is within-mouse variability: blood samples taken from the same animal at the exact same time can result in slightly different fluorescence values. This is due to a varying percentage of plasma present in each blood drop coming out of the punctured site as the blood starts to coagulate. It can be reduced by minimizing the time between puncturing and blood collection and, when sampling weaned mice, by collecting more than one sample from each animal. The second one is inter-mouse variability: blood samples taken from different animals at the exact same time point after injection can vary. The reason is that time zero for injection from birth might differ by a few hours for each batch of pups. These few hours become relevant in how fast Alb-mNG reaches strong expression levels because the liver is growing at a very high rate in the early postnatal phase. Thus, we recommend standardizing the time zero for injection from birth in different studies by closely following the gestation to inject at the beginning rather than the end of PN3. If this is not possible, we recommend sampling the same animals repeatedly in subsequent days until reaching the desired expression levels of Alb-mNG for subsequent procedures.

Note 2.6: The longitudinal study of Alb-mNG expression was stopped at around 15 weeks, i.e. after the expression had reached a constant level. The furthest time point we checked was 6 months from injection in a single animal with comparable expression levels as the 15-week time point (data not shown).

Note 2.7: We chose PN3 as injection day because of our current animal experimentation license. Injection at earlier stages is possible but the injection volume should be adjusted accordingly. Injection at later stages is not recommended for the tested dose, as the liver growth rate is substantially higher in early postnatal periods.

### Note 3: Transcranial imaging

Note 3.1: With our imaging settings, the threshold value for transcranial imaging in pups with transparent skull is 2×10^4^ a.u. whereas for imaging of cranial windows in mice of any age it is 1.5×10^4^ a.u. [10].

### Note 4: Post-mortem validation

Note 4.1: Two- and four-week post-injection time points were chosen due to their peak in Alb-mNG expression and suitability for transcranial imaging.

## Discussion

We present an AAV-mediated CRISPR/Cas9-based approach for knocking in the fluorescent protein gene *mNG* into the albumin locus. A single subcutaneous AAV injection in neonatal mice of 3 days of age lead to successful gene editing in hepatocytes, the predominant producer of albumin. The resulting production of Alb-mNG from genome-edited hepatocytes enabled robust labeling of plasma within a week. Sustained expression was confirmed by *ex vivo* macroscopic fluorescence imaging of blood samples for at least 15 weeks. Because the transgene is stably expressed following chromosomal integration, production of tagged albumin is expected to cover the entire rodent lifespan.

The blood Alb-mNG fluorescence peaked at around 4−5 weeks after injection (Fig. 2b), whereas the knock-in efficiency was higher at 2 weeks post-injection than 4 weeks (Fig. 2g). Between the two postnatal stages, the body weight and total blood volume are doubled. We observed increased total albumin transcript levels at 4 weeks compared to 2 weeks post-injection while *mNG* transcript levels did not differ. This differential transcription profile and thus decremental knock-in efficiency could be explained by genome editing being slower than the liver growth rate, i.e., in these two weeks, the number of total hepatocytes dramatically increases, whereas the pool of genome-edited hepatocytes expands only partially. Nonetheless, considering the plasma albumin concentration of a few hundred micromolar (μM), the estimated 2–6% knock-in efficiency and 1–4 µM Alb-mNG concentration are reasonable.

Compared to our previous method, where a single-shot systemic AAV injection led to episomal expression of Alb-mNG in hepatocytes [10], our knock-in strategy is able to label plasma in the rapidly growing liver of neonate mice. Since the episomal AAV genome dilutes at each cell division, Alb-mNG expression can only be sustained following integration into chromosome 5 of hepatocytes. Another advantage of our knock-in approach is that Alb-mNG is present at physiological expression levels, allowing for a more immune-friendly approach. Indeed, its expression is under the control of the endogenous promoter as opposed to overexpression by AAV8-P3-Alb-mNG. Furthermore, a knocked-in hepatocyte is able to produce labeled albumin without exhausting cell resources for protein synthesis and transport, whereas an overexpressing hepatocyte might be subject to metabolic stress [31]. Finally, our knock-in strategy offers the advantage of producing higher levels of Alb-mNG in both *ex vivo* blood (> 2×10^4^ a.u. in Fig. 2b vs 1.5−2×10^4^ a.u in Wang et al. 2022 [10]) and *post-mortem* plasma samples (1.8−2.8 μM in Fig. 2e vs ∼1 μM in Wang et al. [10]).

A limitation of our method is that injection at early neonatal stages is critical for successful robust labeling of plasma in mice for the tested dosage of 2 × 10^11^ vg. This dosage was validated for injection at PN3 (the earliest day allowed by our current animal experimentation license). We expect that injecting the same dose at PN1 or PN2 is feasible via intravenous injection of volumes up to 50 µL in the temporal vein [32]. Of note, injection of higher volumes (90–100 μL) has been reported [33, 34] but is associated with increased injection-related mortality [32] and could damage the blood brain barrier since the injection volume is comparable to the total blood volume of PN1–PN2 mice [35]. Injection at earlier stages would lead to a faster expression of robust Alb-mNG levels, thus enabling two-photon imaging even before PN10. Given the high liver growth rate in the postnatal period up to 8 weeks of age, injection at stages later than PN3 requires higher dosages. For injection in adult mice, we recommend using AAV8-P3-Alb-mNG [10], as an excessive dosage of the CRISPR/Cas9 AAVs would be required for the fully mature liver. Likewise, larger doses are needed for other animal models according to their liver weight and the relative percentage of hepatocytes. The mouse liver weight is on average 0.1 g at PN3 and 1.4 g at 8 weeks of age [15], whereas the adult liver weight is in the order of 4–9 g in rats, 70–90 g in rabbits and 400–1000 g in pigs [36]. Of note, albumin turnover is much slower in larger animal models: half-life of plasma albumin is 3.6 days in rats [37], 5.7 days in rabbits [18] and 8.2 days in pigs [38], in contrast to 1.2–1.6 days in mice [18, 19].

A straightforward application of our approach is to study brain vasodynamics during development. Cerebral vasculature undergoes consistent expansion and remodeling in the first postnatal month, with the strongest angiogenic activity occurring between PN7 and PN12 [39, 40]. We showed how transcranial two-photon imaging following injection of CRISPR/Cas9 AAVs in PN3 mice enables assessment of vasculature structure, blood flow and vasomotion from PN10. Thinned skull or cranial windows adapted for chronic imaging [41, 42] and/or injection of the same AAV dosage at earlier time points would offer the chance to image neonate mice even before PN10.

Structural and functional changes occur in brain vasculature during aging [43] and in neurodegenerative disorders [44]. Permanent plasma labeling by knocked-in Alb-mNG enables lifelong studies investigating their effects on brain circulation and provides a wide temporal window for evaluating treatment efficacy.

Furthermore, our knock-in strategy can be applied to studying vasculature in peripheral tissue, as we previously demonstrated [10]. Another interesting application could be investigating neurovascular coupling by simultaneous visualization of vasculature, neurons and astrocytes in transgenic mice or in combination with other AAVs (*see* Note 1.11).

This technique holds many promising potentials. The same CRISPR/Cas9 AAVs used in our approach could be directly injected to testis or ovary and the genome-edited sperm or egg cells could be used to create a transgenic mouse line stably expressing Alb-mNG. Furthermore, the CRISPR construct pAAV-U6-AlbEx14-mNG could be engineered with different fluorescent proteins as in our previous paper [10] or bioluminescent proteins. Potentially, albumin could be linked to a therapeutic protein and our CRISPR/Cas9 system could be used for its tissue targeting.

Above all, our CRISPR/Cas9 strategy for persistent labeling of plasma albumin offers a unique opportunity to study cerebral and whole-body vasculature at different spatiotemporal scales from infancy to senescence in the same animal.

## Acknowledgments

This work was supported by NIH BRAIN Initiative U19 Research Program (1U19NS128613-01), Novo Nordisk Foundation (NNFOC0058058), Danmarks Frie Forskningsfond (0134-00107B), JSPS KAKENHI (22K06454, AK; 23K06745 MF), AMED Brain/MINDS (JP21dm0207111, HiH), the Naito Foundation (MasN), and the Lundbeck Foundation. We thank Dan Xue for illustration of experimental schemes. The authors declare no competing financial interests.

## Notes

### Competing Interest Statement

The authors have declared no competing interest.

